# Gut microbiota as a modulator of circadian neural development in the honey bee model

**DOI:** 10.1101/2025.09.30.678393

**Authors:** Yılmaz Berk Koru, Katharina Beer, Angelo Alberto Ruggieri, Josué Alejandro Rodríguez-Cordero, Evelyn Aviles-Rios, Maya Anderson, Esteban A. Citron-Rodriguez, Airined Montes-Mercado, Héctor De Jesύs-Cortés, Manuel Antonio Giannoni-Guzmán, Eddie Perez Claudio, Emma C. Courtney, Cristina Lee Andujar-Sierra, Abigail Strubbe-Nieves, Yarira Ortiz-Alvarado, Mehmet Ali Döke, Humberto Ortiz-Zuazaga, Darrell Moore, Rosanna Giordano, Alfredo Ghezzi-Grau, Ricarda Scheiner, Tugrul Giray, Jose Luis Agosto-Rivera

## Abstract

Disruption in gut microbiota during the early postnatal period can disrupt normal neural development and result in long-term behavioral alterations1. Similar to other neural systems, the circadian clock mechanism continues to mature after birth2, yet how microbial disturbances in the early period influence the onset of circadian rhythms and the development of central clock mechanisms remains poorly understood. Here we studied whether early-life gut dysbiosis affects the ontogeny of behavioral circadian rhythms and the maturation of clock neurons using the honey bee (Apis mellifera), a model organism that shares features of postnatal development of behavioral circadian rhythm and clock system3–5 with humans6. Our findings demonstrate that antibiotic-treated and gnotobiotic-reared bees display reduced rhythmicity compared to controls. These treatments also impair the development of the circadian pacemaker, marked by fewer Pigment-Dispersing Factor (PDF)-expressing neurons. Additionally, antibiotic exposure increased the expression of the Insulin-like Growth Factor Binding Protein Acid Labile Subunit (IGFALS) in early ages, which stabilizes the IGF-1/27, a hormone important for neurodevelopmental processes42. Together, these results identify gut microbiota as a modulator of circadian development. Our work provides an understanding of how early-life microbial disruptions influence the development of circadian rhythms, providing information that may extend to other animals, including humans.

## Main

Exposure to adverse environmental factors during the postnatal period can profoundly influence neural development, plasticity, and subsequent behavior^8^. One aspect of postnatal brain development is the circadian system, which continues maturing after birth in mammals, or eclosion in eusocial insects including honey bees^5,9,10^. Circadian rhythms are daily oscillations in physiology and behavior that synchronize to anticipate environmental cues^11^. Disruption of circadian rhythms by factors such as irregular light exposure, environmental toxins, and unscheduled feeding has been associated with a variety of health consequences^12–15^. Also, disruption of the gut microbiota alters circadian gene expression in the liver, gut, and brain, which in turn affects host metabolism and behavior^16–18^. Yet, the impact of these adverse environmental factors on the development of circadian rhythms remains poorly understood^2^.

A growing body of evidence has shown that gut microbiota plays a role in brain development and behavioral outcomes^1^. The gut microbial community is highly sensitive to environmental perturbations, with antibiotic exposure capable of inducing dysbiosis, which is a disruption in microbial balance^19^. Recent studies conducted on rodent models show that depletion of gut microbiota through either antibiotic treatment or in germ-free organisms has been linked to altered microglial activity and synaptic function, changes in astrocyte function, dysregulated gene expression as well as behaviors including learning, social interaction, and cognition^20–27^.

With this study, we addressed if early postnatal-period microbial disruption influences the development of behavioral circadian rhythms and the clock system of the honey bee (*Apis mellifera*). model system that allows the manipulation of gut microbiota through antibiotic treatment and gnotobiotic rearing^28^. Honey bees also exhibit post-embryonic development of behavioral circadian rhythms and clock systems similar to humans^3–5,29–31^ permitting free-running behavioral assays under controlled conditions. We hypothesize that gut microbes are essential for the ontogeny of the behavioral circadian rhythm and the maturation of the neural clock. We assess how microbial disruption affects the development of circadian rhythms using locomotor activity assays to evaluate behavioral rhythmicity; immunostaining to quantify the number of pigment-dispersing factor (PDF)-expressing neurons in the brain; rRNA-depleted RNA-Seq of gut tissue to examine the impact of antibiotic treatment on the microbial flora; and qRT-PCR of brain tissue to measure the expression of *IGFALS* across ages.

### Gut microbiota disruption impacts the development of behavioral circadian rhythm

Worker honey bees undergo post-embryonic brain and behavioral development^32,33^. Initially, workers perform cell-cleaning, then transition to nursing, and later perform other hive duties such as ventilation, defense, and food processing. Circa three weeks after emerging as adults, bees begin foraging and gather nectar, pollen, and water^34^. However, this behavioral development can be influenced by chemical stressors, including antibiotics^35^. Ortiz-Alvarado et al. (2020) reported delayed behavioral development in honey bees exposed to antibiotics, and fewer foragers and more cleaning behaviors observed between days 11 and 14 post-emergence. Behavioral development is tightly associated with the onset of circadian rhythmicity in honey bees^3,36^.

To examine whether the antibiotic Tylosin Tartrate affects the development of behavioral circadian rhythms, newly emerged bees were exposed to the antibiotic for 14 days and individual locomotor activity monitored using Locomotor Activity Monitors (LAMs). We hypothesized that antibiotic-induced dysbiosis would impact the onset of behavioral rhythmicity. We observed that by day 4, the antibiotic treatment group had 4.87% rhythmic individuals, whereas the control group had 13.88%. This trend persisted throughout the experiment, with 41.46% of treated bees versus 63.88% of controls showing rhythmic behavior by day 14 (Fig. 1B). We next compared circadian parameters (period, rhythm strength) among rhythmic individuals during the last three days of the assay. No significant differences were observed in either period or rhythm strength (Supplementary Fig. 1A–B). Mortality rates were also similar between groups (Supplementary Fig. 1C).

**Figure 1.**
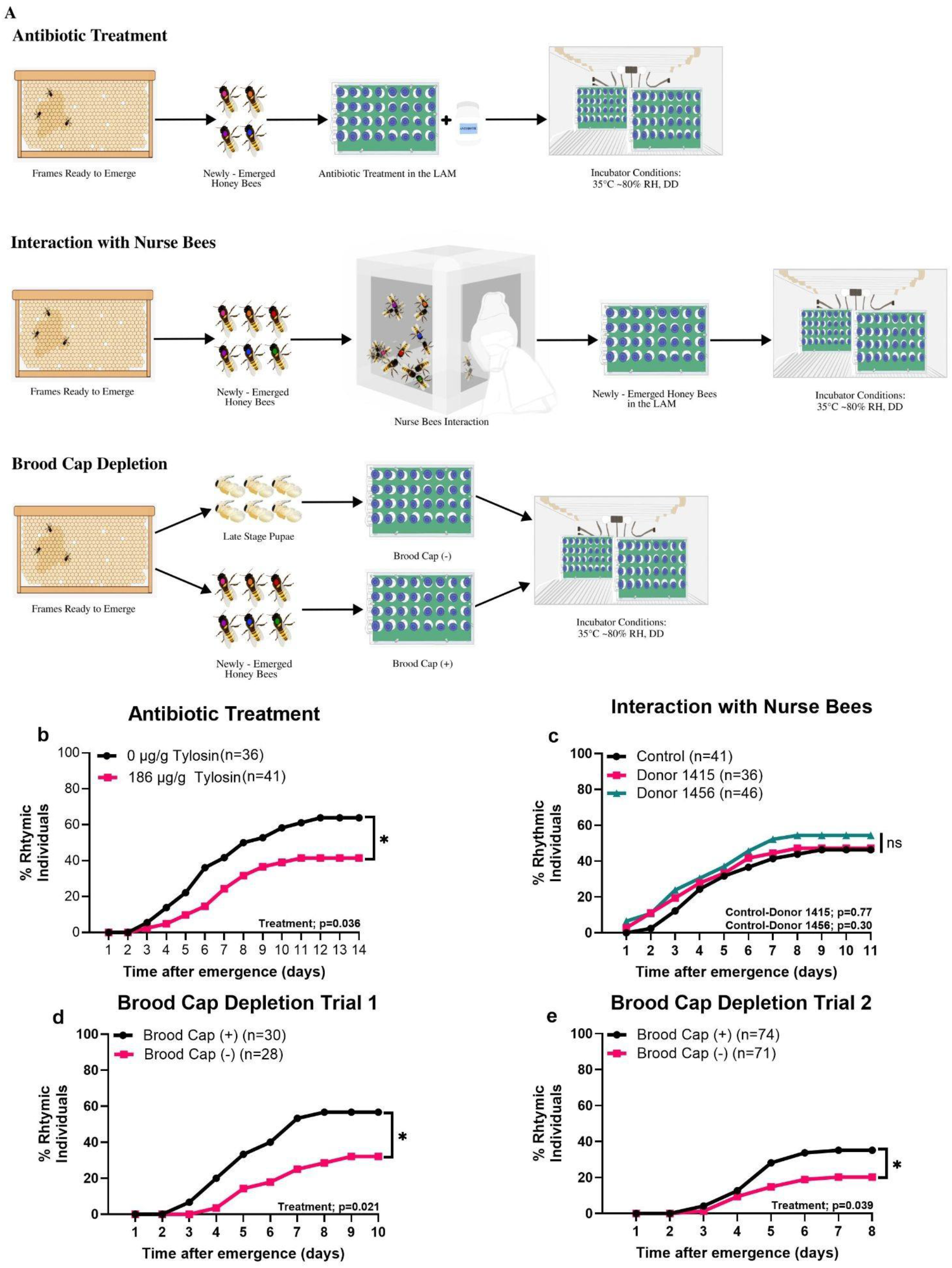
**A**, Manipulation applied for the behavioral assay; **B**, Antibiotic treatment on the ontogeny of behavioral circadian rhythm (Wald = 4.39, p = 0.0361, General Estimating Equations) **C**, Interaction with nurse bees on onset of behavioral development (Donor 1415; Wald test = 0.09, p = 0.77, Donor 1456; Wald test = 1.09, p = 0.30, Generalized Estimating Equations) **D**,**E** Brood cap depletion on ontogeny of circadian rhythm (Brood Cap Depletion Trial 1: Wald test = 5.3, p = 0.021, Generalized Estimated Equations; Brood Cap Depletion Trial 2: Wald test = 4.24, p = 0.039, Generalized Estimated Equations).

To account for a potential microbiota-independent pathway by which antibiotics may adversely influence host behavior^37^, we employed an alternative microbial manipulation: newly emerged bees were exposed to nurse bees from two different colonies (Donor 1415 and Donor 1456), creating two independent donor groups to ensure robust microbial transmission and to control for colony-specific effects. In each treatment group, nurse bees transferred microbiota through oral trophallaxis to newly emerged bees while the control group bees were not exposed to nurse bee interaction. Our results showed no significant differences across groups. All groups i.e. control, donor 1415, and donor 1456 exhibited comparable rhythmicity levels: 46.34%, 47.22%, and 54.34%, respectively (Fig. 1C).

As a third approach, we employed a brood cap depletion method by removing caps from brood cells. Naturally emerging bees chew through the brood cap, potentially acquiring gut microbes during this process^28,38^. We removed late-stage pupae from their cells before eclosion, resulting in no or reduced microbial exposure from the brood cap^39^. Data from this experiment was used to assess if microbes acquired from the brood cap influence the development of circadian rhythms. In two trials, we observed that bees that emerged naturally (brood cap (+)), Trial 1, developed behavioral rhythms at higher rates than those in Trial 2, removed before eclosion (brood cap (−)) (56.66% and 32.14%, Trial 1; 35.21% and 20.27%, Trial 2) (Fig 1D-E). There were no significant differences in rhythm strength, period, or survival (Supplementary Fig 3 A-D). These findings suggest that early microbial exposure via the brood cap supports behavioral circadian development.

Taken together, our findings show that gut microbiota dysbiosis impairs the development of behavioral circadian rhythms, as demonstrated by both antibiotic treatment and brood cap microbiota depletion. In contrast, microbial transfer, or lack thereof, through nurse bee interaction did not have a significant effect.

### Antibiotic treatment altered gut microbiota composition

To evaluate the impact of Tylosin Tartrate on honey bee gut microbiota, we performed rRNA depleted RNA-Seq on gut tissues from 14-day-old bees subjected to a dose-response experiment. Microbial composition shifted significantly at family level (PERMANOVA: F(1,9) = 1.81, P = 0.0478) (Supplementary Figure 4A). Antibiotic treatment reduced the relative abundance of core bacterial families, including Neisseriaceae, Bifidobacteriaceae, Lactobacillaceae, and Orbaceae, as well as Acetobacteraceae and Pseudomonadaceae. In contrast, the relative abundance of Enterobacteriaceae and Yersiniaceae increased, both of which include opportunistic bacterial species (Motta & Moran, 2024). Additionally, even though the antibiotic treatment caused reduction in the alpha diversity around the 40.68% of mean of Shannon index, there was no significant differences (Supplementary Figure 4B).

**Figure 2.**
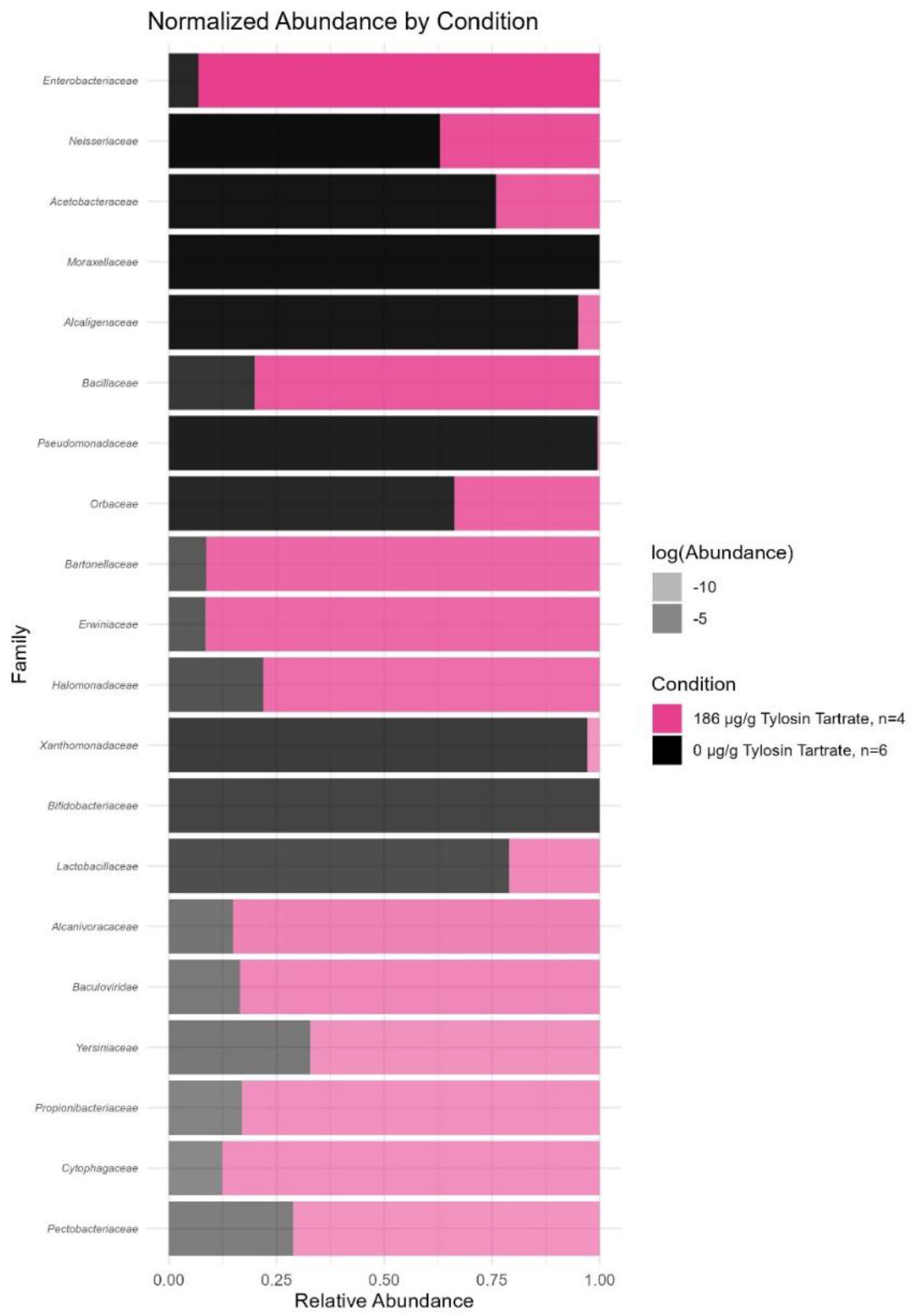
the effect of antibiotics on gut microbiota composition, focusing on the 20 bacterial family with the highest relative abundance.

### Dysbiosis in gut microbiota affects maturation of Pigment-Dispersing Factor-expressing neurons

Pigment-Dispersing Factor (PDF), a neuropeptide influencing locomotor activity in honey bees, is produced by clock cells that also express the central clock protein Period (Per)^10,41^. Beer and Helfrich-Förster (2020) demonstrated that young, arrhythmic honey bees exhibit significantly fewer PDF-expressing neurons than older, rhythmically active bees.

To examine whether gut microbiota contribute to the development of the neural clock system, we performed immunostaining of PDF-expressing neurons following two microbiota manipulation approaches: antibiotic treatment and brood cap depletion. Our findings revealed that bees treated with antibiotics exhibited a significantly lower number of PDF-expressing neurons compared to control bees (Fig. 3A). In the brood cap assay, bees that emerged after chewing through the brood cap had more PDF-expressing neurons than those that did not (Fig. 3B). Notably, rhythmicity had no significant effect within the brood cap (+) group. However, in the brood cap (−) group, arrhythmic bees exhibited significantly fewer PDF-expressing cells compared to rhythmic individuals. Our results show that dysbiosis in the gut microbiota by either antibiotic treatment or brood cap deprivation impair PDF neuron maturation.

**Figure 3.**
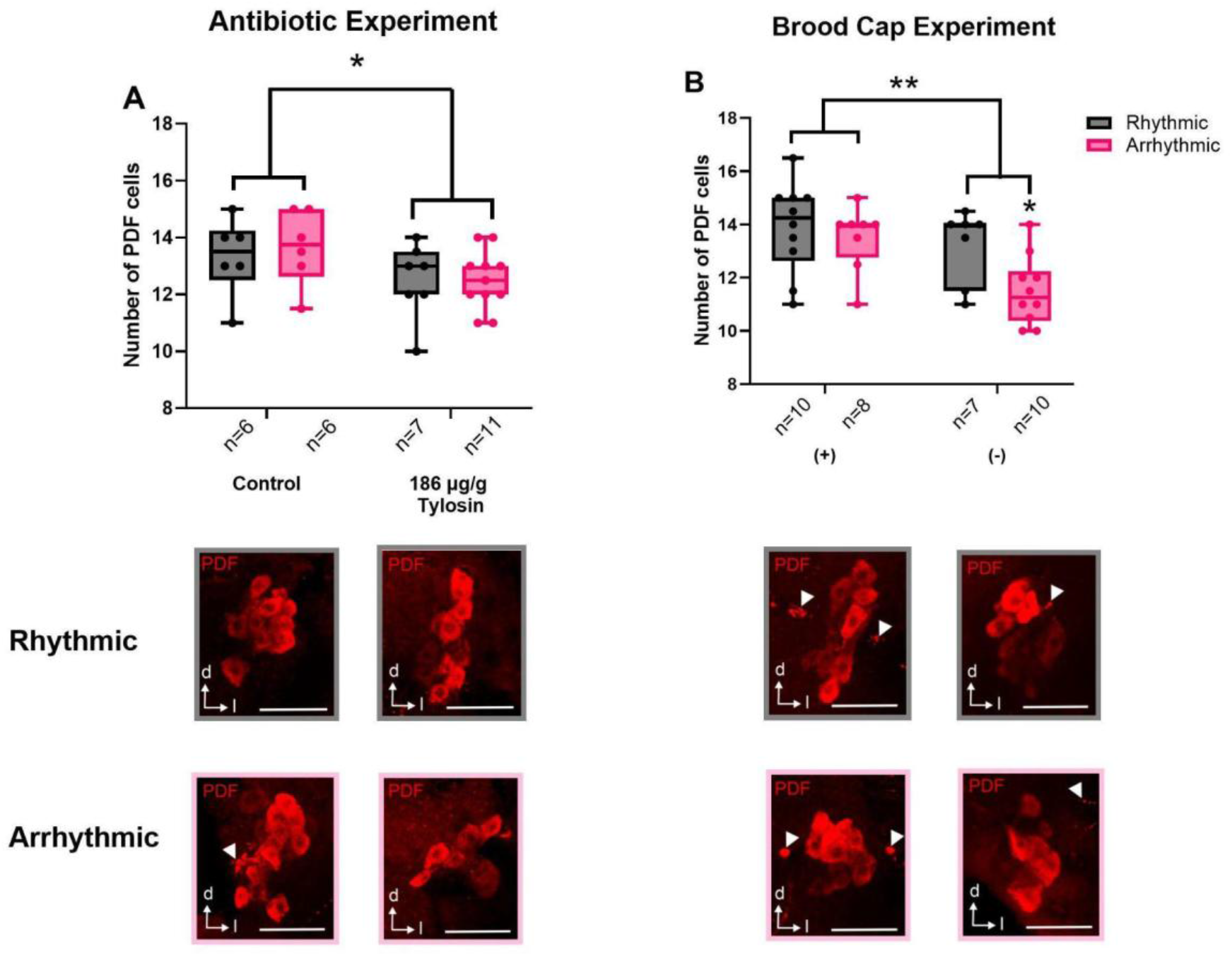
**A**, Antibiotic treatment effect on PDF expressing neurons (Interaction: F(1, 26) = 0.1301, P = 0.7228; Treatment effect: F(1, 26) = 4.7796, P = 0.03799; Rhythmicity effect: F(1, 26) = 0.0912, P = 0.76504, Generalized Linear Model) **B**, Brood cap depletion effect on PDF expressing neurons (Interaction: F(1, 31) = 1.8217, P = 0.186888; Treatment effect: F(1, 31) = 9.9186, P = 0.003608; Rhythmicity effect: F(1, 31) = 4.4998, P = 0.041998; Generalized Linear Model). Arrows on the representative images point out fibers. d= dorsal, l= lateral. Scale bar: 50 μm, values and error bars mean ± s.e.m.

### Antibiotic treatment affects the expression of a neural development gene

To investigate the impact of antibiotic treatment on gene expression, we first performed a preliminary RNA-seq analysis on whole brain tissue. We focused on genes showing a significant interaction between antibiotic treatment and rhythmicity. Differential expression analysis was conducted using two packages: DESeq2 and Limma. DESeq2 identified 50 differentially expressed genes (Supplementary List 1A), while Limma identified 62 (Supplementary List 1B). We then selected the genes common to both analyses, resulting in 32 shared genes (Supplementary List 1C; p-value < 0.05, |log2 fold change| > 1).

Among these, one notable gene was *insulin-like growth factor-binding protein acid labile subunit* (*IGFALS*). *IGFALS* forms ternary complexes with IGF-1/2 and IGF-binding proteins (IGFBP-3 and IGFBP-5), stabilizing IGFs in the circulatory system by prolonging their half-life and modulating their bioavailability^7^. IGF signaling plays a critical role in neural development^42^. To evaluate the impact of antibiotic treatment on *IGFALS* expression across different developmental stages, we performed absolute qRT-PCR. Expression levels were quantified at four specific post-eclosion time points: days 2, 4, 6, and 8.

Our results show that there is a significant difference between expression across treatment groups over time. Specifically, on day 2, the antibiotic-treated group exhibited a significantly higher copy number of *IGFALS* transcripts compared to the control. These findings suggest that antibiotic-induced dysbiosis upregulates *IGFALS* expression during early post-emergence, possibly impacting neuronal development via modulation of *IGF* signaling.

**Figure 4.**
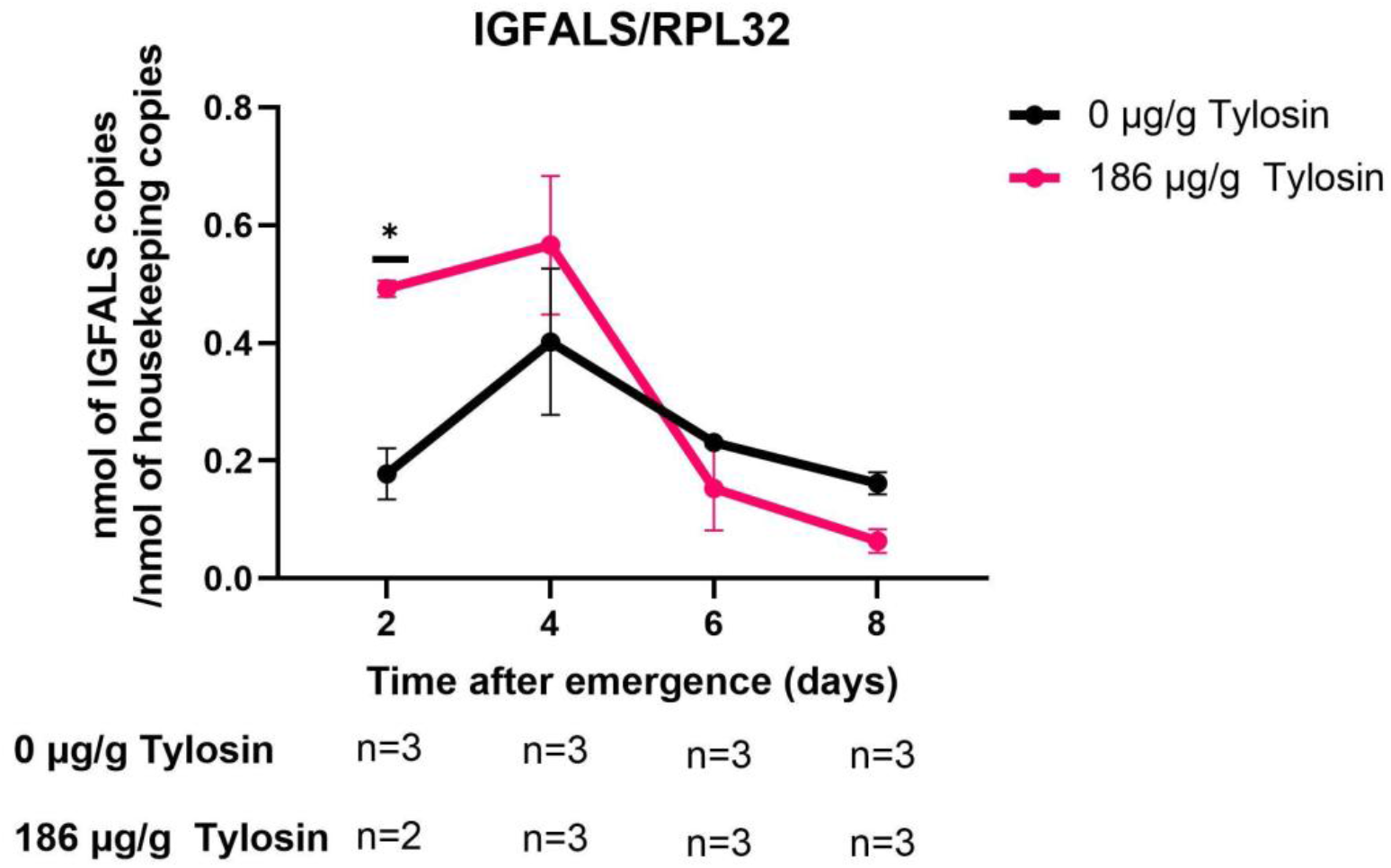
Antibiotic treatment on the IGFALS expression in the honey bee brain across developmental time points (Interaction: F(3,15) = 3.5962, P = 0.03878; Days: F(3,15) = 10.7469, P = 0.0005; Treatment: F(1,15) = 2.1807, P = 0.16043, Generalized Linear Model); Day 2: P value = 0.0127; Day 4: P value = 0.1191; Day 6: P value = 0.4448; Day 8: P value = 0.3398, values and error bars mean ± s.e.m).

## Discussion

Here, we show that in honey bees, gut microbiota plays a role in the development of the behavioral circadian rhythm and maturation of the clock system. Honey bees exposed to early-life gut microbiota disruptions, through antibiotic treatment or brood cap depletion, exhibited significantly reduced behavioral circadian rhythmicity. Newly emerged bees that interacted with nurse bees did not show significant differences in the development of circadian rhythm of locomotor behavior compared to controls which did not interact with nurse bees, suggesting that microbial transfer through social contact does not influence circadian development at the behavioral level.

Furthermore, the antibiotic treatment and brood cap depletion experimental manipulations, which resulted in the disruption of gut microbiota significantly impaired the maturation of Pigment-Dispersing Factor (PDF)-expressing neurons, the pacemaker cells of the brain. Our findings also demonstrate that microbiota disruption affects not only the behavioral outcome of the bee circadian rhythm but also neurodevelopmental pathways that may influence circadian maturation. Specifically, antibiotic exposure led to increased expression of IGFALS at early ages. Gut microbiota may help regulate IGF ligand bioavailability or the dynamics of complex dissociation^27^. In honey bees exposed to antibiotic treatment, disrupted microbial signaling may lead to the accumulation of unbound IGFALS, reducing the availability of IGFs for receptor binding in clock neurons. This could impair neuronal maturation and disrupt the development of behavioral circadian rhythms.

Previous studies have shown that dysbiosis of the gut microbiota impaired the host’s circadian rhythm gene expression in the liver, intestine, and brain in mice^16–18^. Our findings demonstrate for the first time that dysbiosis of gut microbiota in the honey bee impairs the development of behavioral circadian rhythm and the clock system. Future work should characterize specific bacterial species and their metabolites involved in this process. In summary, our study establishes gut microbiota as a modulator of circadian neurodevelopment in honey bees, a finding potentially applicable to mammals, including humans.

## Material and Methods

### Honey Bee Collection

Honey bee colonies used in this experiment are maintained with standard methods at the Gurabo Experimental Station apiary at the University of Puerto Rico, Rio Piedras. Brood frames ready to emerge were picked from healthy colonies to obtain both pupae and newly-emerged worker bees. Pupae were collected on the same day that frames were collected. Frames used to collect newly-emerged bees were placed overnight in an incubator (Percival Intellus) set at 35°C and 80% ± 5% relative humidity. The antibiotic treatment experiments, including behavior, gene expression and immunostaining assays, were started on February 4, April 19, 2021, March 14, 2022 and May 23, 2023. Experiments 1 and 2 for microbe transfer by nurse bees took place on July 27 and September 14, 2022, respectively. The brood cap experiments, which also involved behavior and immunostaining assays, were performed on March 1 and June 8, 2023.

### Locomotor Activity Monitors

Locomotor Activity Monitors (LAMs), manufactured by Trikinetics Inc. (Waltham, MA, USA), were used to measure the movement of the honey bees. Each monitor consists of 32 channels, each incorporating three infrared beams for detecting locomotor activity of individual honey bees housed in 15 ml Eppendorf tubes. All monitors were prepared as previously described by Giannoni-Guzmán et al. (2014). Behavior and gene expression experiments were conducted inside an incubator set at 35°C, with a relative humidity of 80%±5%, under constant darkness for only behavior and gene expression experiments2.

While the honey bees were in the LAMs, food and water were provided “ad libitum”. Water was refilled every three days, and the food supply was checked one week later to assess its sufficiency. The honey bees in the antibiotic treatment and nurse microbe transfer experiments were fed with honey candy, which is prepared with honey and powdered sugar at a ratio of 1:3.6 (honey/powdered sugar). Honey bees in the brood cap assay were fed with ground sucrose (Sigma-S7903) mixed with 2M sucrose solution at a ratio of 1:5.9 (sucrose solution/ground sucrose), and food used in both brood cap experiments was prepared inside a class II biological safety cabinet, minimizing microbial contamination. Honey bees from multiple colonies were populated on the LAMs in a specific order so all colonies were included equally or approximate numbers.

### Antibiotic Treatment Experiments

Eight newly emerged bees from four different colonies were placed in each LAM for both the antibiotic treatment and control groups. The newly emerged bees in the antibiotic treatment group were exposed to the antibiotic Tylosin Tartrate (Tylan, Elanco, Product Label: AF1300) added to their food at a 186 µg/g dose for 14 days until the end of the experiment. The concentration of the antibiotic used was determined via a dose-response curve measuring the magnitude of the effect in rhythmicity if there was a delay on the development of locomotor circadian rhythm.

### Microbe Transfer by Nurse Bees Experiments

To investigate the role of nurse bees in microbiota transfer, we allowed them to interact with newly emerged bees in controlled cage experiments, following the approach of Powell et al. (2014). Frames with brood ready to eclose were taken from six colonies and placed in an incubator overnight. The next day, newly emerged bees were marked on their thoraxes with one of six different colors to indicate their colony of origin. Nurse bees were identified as individuals inserting their heads into cells containing larvae.

In the Trial 1 (Supplementary Fig 2 A-E), we collected approximately 300 nurse bees, half from donor colony 1460 and half from donor colony 1468. These were housed separately in two BugDorm-1 insect rearing cages (30 × 30 × 30 cm) with mash panels on the back, left, and right sides, and clear panels on the bottom, front, and top cages based on their colony of origin. Additionally, three groups of 200 newly emerged bees were collected with equal representation (∼33 bees) from six different colonies and marked with unique colors for identification. Each group of 200 bees was placed in the cages. Two groups were housed with nurse bees, while a third group served as a control without nurse bees. However, nurse bees attacked the newly emerged bees, leading to reduced sample sizes and altered mortality trends compared to other experiments.

To address these issues, we refined the experimental design by increasing the sample size and sourcing nurses and newly emerged bees from different colonies. We collected 400 nurse bees, half from donor colony 1415 and half from donor colony 1456. Three groups of 300 newly emerged bees, each comprising ∼50 individuals from six different colonies, were introduced into two experimental cages and one control cage without nurse bees. During both the Trial 1 and Trial 2, newly emerged bees were kept with nurse bees in cages for two hours before being transferred to individual tubes in Locomotor Activity Monitors (LAMs).

### Brood Cap Experiments

Brood frames were collected from six different colonies. On the same day that frames were collected, pupae were taken out of the frames as described in Powell et al., 2014, placed in tubes inside the LAMs, and eclosed without chewing the brood cap. This method aimed to minimize bacterial load3,4, as previous studies have suggested that emerging bees may acquire microbes by chewing the brood cap during the eclosion including alpha, gamma, and firm-5 phylotypes5. The following day, newly emerged bees that chew the brood cap were collected from the frames and placed in the LAMs.

### Immunostaining of PDF-Expressing Neurons in Honey Bees

#### Sampling for Immunostaining

Two different experiments were set for the immunostaining assay (immunohistochemistry, IHC): antibiotic treatment and brood cap depletion. For the brood cap depletion experiment, six frames of brood ready to emerge were collected. As different from the protocol mentioned in the brood cap experiment section, pupae, and newly emerged bees were placed in the LAMs on the same day.

For the antibiotic experiment, five brood frames were ready to emerge and be picked. The same protocol was used as mentioned in the antibiotic experiment to rear the honey bees and measure their locomotor activity.

The constant dark condition for 8 days was then changed to a light-dark cycle (LD 12:12 h, with lights on at 9:00 and lights off at 21:00 h) in the last three days to synchronize individual clocks of the honey bees and thereby establish standardized conditions for the PDF-expressing cell quantification across treatments.

For the IHC experiments involving the brood cap, honey bees were classified into different groups based on two criteria: chewing the brood cap and their rhythmic behavior. Similarly, for the IHC experiments regarding antibiotic treatment, honey bees were divided into groups according to two factors: the administration of antibiotic treatment and their rhythmic behavior.

#### Immunostaining

After retrieving the honey bees from the LAMs, honey bees were anesthetized using regular ice, decapitated, and part of the cuticle was removed to open a window using a blade. The heads were fixed using Zamboni’s fixative (Newcomer Supply part #1459A) to preserve and stabilize brain tissue and placed overnight at 4°C in a tube rotator at a low speed similar to Beer and Helfrich-Förster (2020). After fixation, the heads were washed in PBS 1x until the solution was transparent with each wash lasting 10 minutes. Brains were dissected in phosphate-buffered saline (PBS) 1x and washed three times in PBS 1x after the dissection. For storage at -20°C, the solution underwent a sequential change in methanol concentrations within 1x PBS, progressively increasing through stages of 30%, 50%, 70%, 90%, and 100% (each stage 10 minutes on a shaker). On the day before shipping to the University of Würzburg/Germany, where the immunostaining assays were performed, the brains were rehydrated with methanol/PBS 1x concentrations (90%, 80%, 70%, 50%, and 30%, 10 mins on a shaker each) and washed three times for 5 mins in PBS 1x on a shaker at room temperature. Brains were shipped in 70% ethanol/PBS 1x (30%, 50%, 70% 10 mins on a shaker for each washing). Before immunostaining, brains were rehydrated in reverse concentration on the same conditions and then washed in PBS 1x three times for 5 mins on a shaker.

The DEpigmEntation-Plus-Clearing method (DEEP-Clear) was used to prepare brains, with modifications to adapt the protocol for the smaller honey bee brain. This method, originally developed for use in worms, squid, axolotls, and zebrafish7, was adjusted as follows. The brains were placed in 100% acetone and incubated at -20°C for 30 minutes, followed by washing with PBS 1x five times for 20 minutes each at room temperature. After this step, the brains were washed with glycine (2 mg/ml) two times for 10 minutes at room temperature, and afterward were washed five times for 20 mins on the shaker using PBS 1x at room temperature. Brains were placed in a solution of 10% (v/v) THEED, 5% (v/v) Triton X-100, and 5% (w/v) urea in dH2O (solution 1.1) for 90 minutes in a water bath at 37°C, and then washed with PBS 1x, five times for 20 minutes.

For immunostaining, the brains were washed three times in PBS 1x, one time in PBST (0.5% Triton), one time in PBST (2% Triton), and one time in Phosphate Buffered Saline-Triton (PBST) (0.5% Triton) for 10 minutes of each on a shaker. The brains were incubated in 5% normal goat serum (NGS)-PBST (0.5% Triton) overnight at 4°C on a shaker. The brains were placed in a primary antibody solution consisting of both 1:3000 anti-β -Pigment Dispersing Hormone (PDH) which recognizes PDF in various organisms, including honey bees8,9, and PBST (0.5% Triton), 5% NGS, and 0.02% NaN3 at 4°C for six days and at room temperature for one day, then washed six times with PBST (0.5% Triton) for 10 minutes each on a shaker. Afterward, the brains were treated with a secondary antibody solution, 1:400 Alexa anti-rabbit 633 antibody for PDH. PBST (0.5% Triton) and 5% NGS at 4°C on the shaker overnight, covered with aluminum foil to prevent light degradation. Brains were washed three times in PBST (0.5% Triton) and three times in PBS for 10 minutes each on a shaker at room temperature.

Silicone spacers were used to mount the brains, which were placed on glass slides (22mm × 22mm × 0.7mm) with antifade mounting media (Vectashield, Vector Laboratories) and covered with a glass coverslip. A Leica SP8 (DM6000) confocal microscope was used to image the cell bodies (University of Würzburg in Germany). All brains were scanned with 20x/0.75 IMM HC PL APO CS2 objective, and the same settings, which were 1024 × 1024 resolution, were used for both experiments. Cell bodies were counted, and the average for images containing two hemispheres. For the images having one hemisphere, the number of stained cell bodies was used.

### RNA-Seq Assay from Gut Tissue

#### Sampling for the RNA-Seq Assay

Honey bee samples were collected from a dose-response experiment in which bees of the same age as those in the behavioral experiment were exposed to the same antibiotic concentration. On the final day of the experiment (day 14), the bees were removed from the LAMs, euthanized by flash-freezing, and their abdomens were separated on dry ice. The abdomens were placed in 750 µL of RNAlater-ICE (Invitrogen AM7030) and stored at -20 °C for approximately two days to penetrate RNAlater-ICE to the tissue.

Gut dissections of the abdomens were done on a glass surface. Guts were carefully removed using tweezers by holding the bee stingers and placed in 750 µL of RNAlater-ICE until RNA extraction.

#### RNA Extraction and RNA-Sequencing Assay

RNA from gut tissues was extracted using the EZ1 RNA Tissue Mini Kit and quantified with the Agilent 2100 Bioanalyzer. rRNA depletion method was performed using the Illumina Ribo-Zero Plus rRNA Depletion Kit (Catalog ID: 20037135), following manufacturer guidelines with following modifications. Library preparation was conducted on a sciclone liquid handler using NEXTFLEX-Rapid-Directional-RNA-Seq-Kit (Catalog ID: NOVA-5190-02) and then sequenced using an Illumina Novaseq 6000 S1 system and 50 base paired end created. For data processing, the quality of sequencing from each sample was assessed using FASTQC/0.11.9. Low-quality sequences were removed using Trimmomatic 0.3910. To remove the honey bee RNA, we used Bowtie2 with very-sensitive and –un-conc-gz settings. After this step, Kraken2 was used to determine the bacterial taxonomy with confidence threshold of 0.1.

#### Absolute Quantification qPCR Assay for Insulin-like Growth Factor-Binding Protein Acid Labile Subunit from Brain Tissue

The gene Insulin-like Growth Factor-Binding Protein Acid Labile Subunit (IGFALS) was identified in honey bee brain tissue via preliminary RNA-Seq. It was common to the differentially expressed gene (DEG) lists generated by both DESeq2 and Limma.

Honey bees from both control (0 µg/g Tylosin Tartrate) and antibiotic-treated groups (186 µg/g Tylosin Tartrate) were reared in LAMs under constant darkness (DD) conditions. Samples were collected every two days (days 2, 4, 6, and 8), and bees were sacrificed by flash-freezing in liquid nitrogen, followed by placement on dry ice to prevent RNA degradation. Samples were stored at –80 °C until brain dissection, which was performed on dry ice. Dissected brains were transferred to 300 µL of RNAlater-ICE (Cat. No: Invitrogen AM7030) and stored at –20 °C until RNA extraction.

Brain tissues were homogenized using a QIAGEN TissueLyser II, and RNA was extracted using the RNeasy Mini Kit (QIAGEN, Cat. No: 74106) on the QIAcube Connect. RNA concentration and purity were assessed using 2 µL of each sample on a QIAxpert system. Complementary DNA (cDNA) synthesis was conducted with Bio-Rad iScript Reverse Transcription Supermix (Cat. No: 1708841).

To prepare a dilution series for absolute quantification, 1 µL of cDNA from each sample was pooled into a single tube, and the genes *IGFALS* and the housekeeping gene *RPL 32* (Supplementary List. 1) were amplified using OneTaq 2X Master Mix with Standard Buffer (NEB, Cat. No: M0482) on a Bio-Rad DNA Engine thermal cycler under the following conditions: initial denaturation at 94 °C for 30 sec, followed by 30 cycles of denaturation at 94 °C for 20 sec, annealing at 57 °C for 20 sec, extension at 68 °C for 30 sec, and a final extension at 68 °C for 5 min. PCR products were purified using the QIAquick PCR Purification Kit (QIAGEN, Cat. No: 28106).

Absolute quantification qPCR was then carried out using the QIAGEN Rotor-Gene Q system with the QuantiNova SYBR Green PCR Kit, following the manufacturer’s protocol. Each 10 µL reaction mixture included 0.5 µL cDNA, 5 µL SYBR Green Master Mix, 0.7 µL each of 10 µM forward and reverse primers, and 3.1 µL nuclease-free water. The qPCR thermal profile consisted of an initial denaturation at 95 °C for 2 min, followed by 40 cycles of 95 °C for 5 sec and 60 °C for 10 sec, with a final melting curve step starting at 65 °C for 1 min 30 sec.

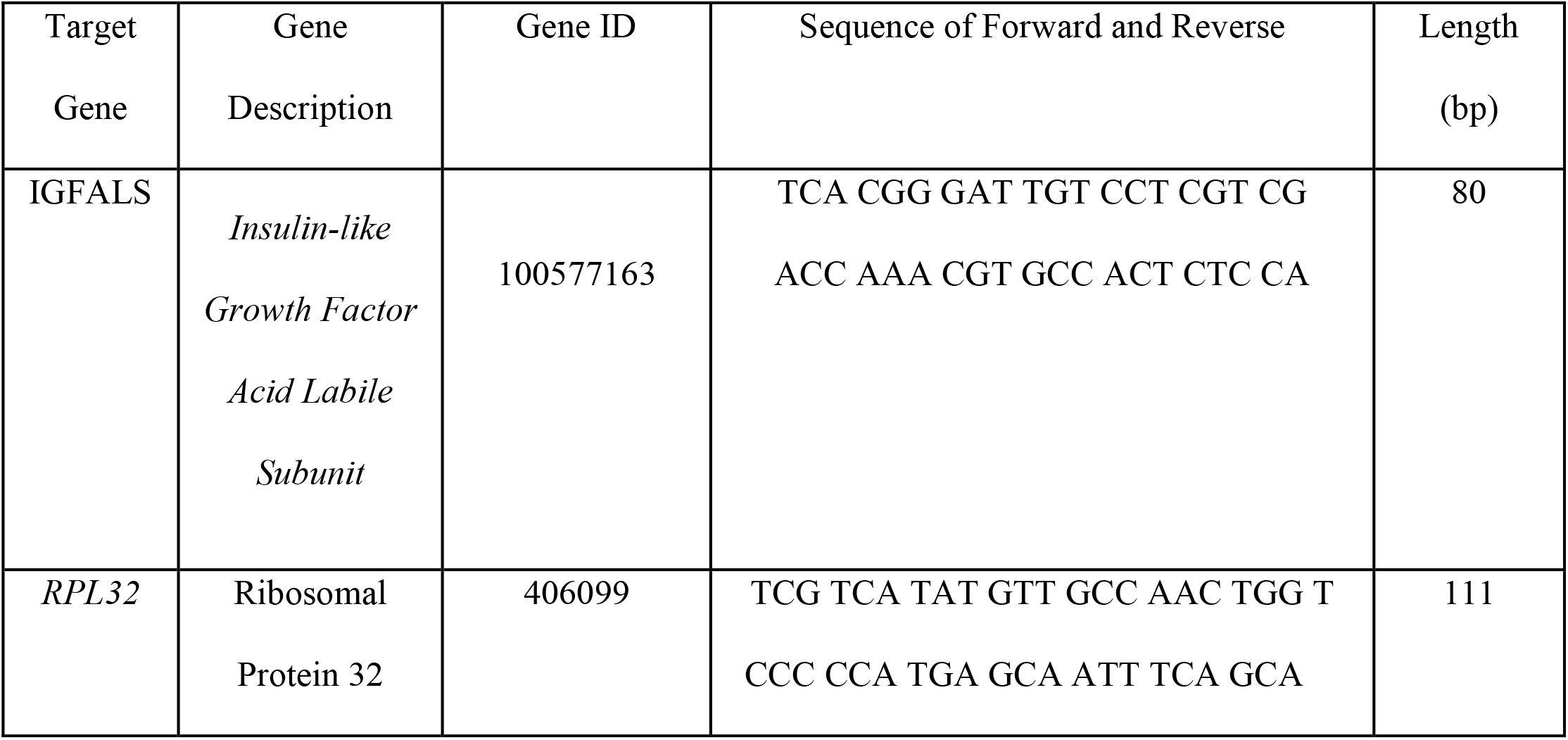

#### Experimental Design and Statistical Analyses

The rhythmicity and onset of activity in individual bees were assessed using the Lomb-Scargle Granger test, combined with visual inspection of double-plotted actogram for each bee (Figures 1-3). For visual inspection, we identified a rhythm based on consistent movement and rest patterns occurring at approximately the same time each day. Also, we obtained period and rhythm strength from lomb-scargle analyses. Survival was evaluated using a double-plotted actogram. Worker bees were considered dead when prolonged inactivity was observed in the tube, with death visually confirmed afterward. To statistically assess group differences in the onset of rhythmicity, Generalized Estimating Equations (GEE) models were applied^12^. Period and rhythm strength were compared between the two groups using a Generalized Linear Mixed Model (GLMM), with colony included as a random effect in all three experiments. For the nurse bee interaction and brood-cap experiments, trial was also included as an additional random effect. Survival analyses were performed using the Kaplan–Meier method and the log-rank test. Additionally, Generalized Linear Model (GLM) was conducted to evaluate group differences in immunohistochemical staining and absolute qPCR quantification assays. Bacterial community structure divergence between control and antibiotic treatment was assessed using Permutational Multivariate Analysis of Variance (PERMANOVA). Normality was tested with the Kolmogorov–Smirnov test. Actogram visualizations and Lomb-Scargle analyses were generated using the Circadian Dynamics R package (https://github.com/edpclau/circadian-dynamics). All statistical computations were performed using R and GraphPad Prism version 10 (GraphPad Software, La Jolla, CA, USA) and R software, while all graphical representations were created in GraphPad Prism version 10 (GraphPad Software, La Jolla, CA, USA) and R software.

## Supplementary Figures

### Antibiotic Treatment Experiment

**Figure 1:**
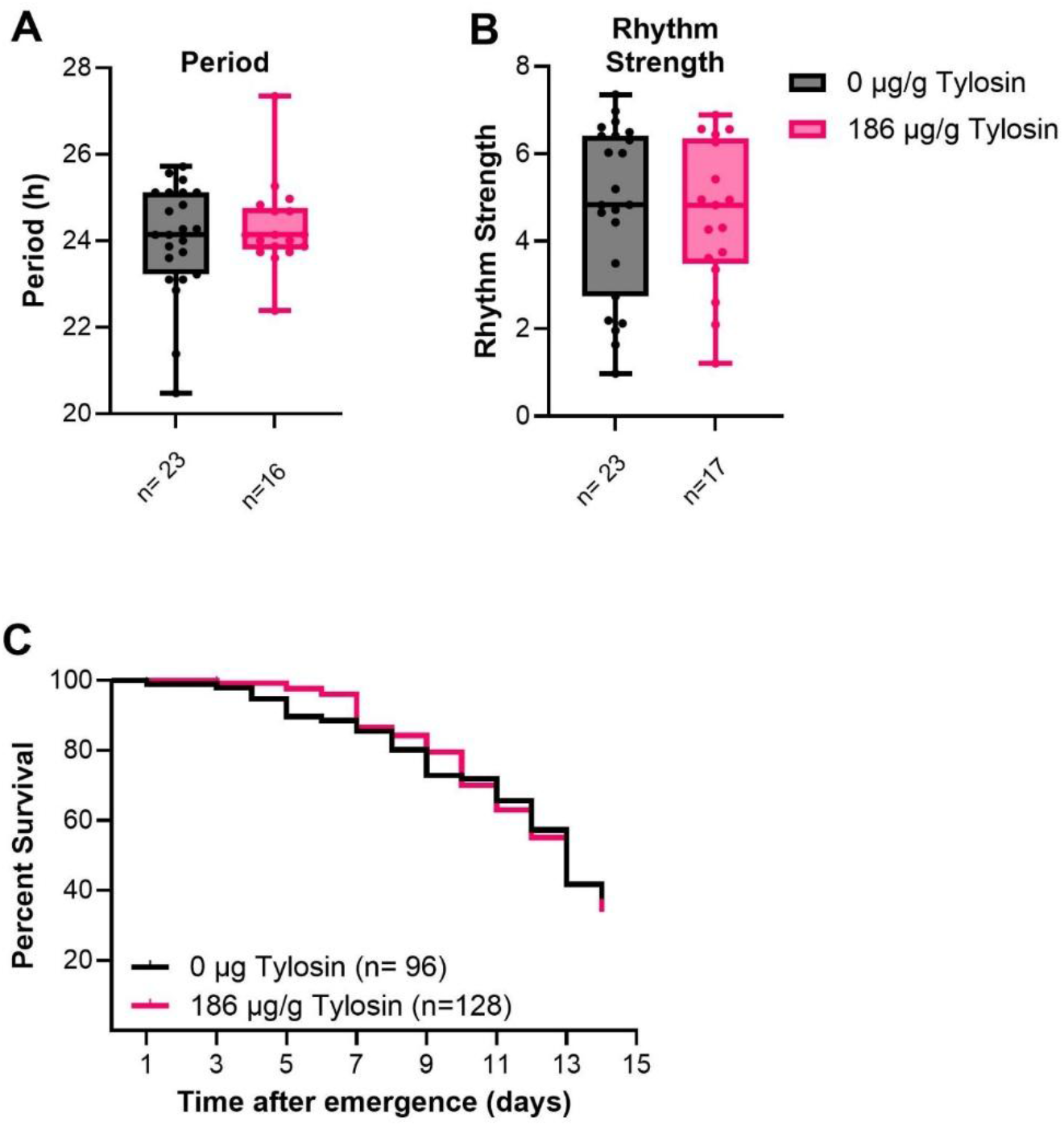
Antibiotic treatment did not affect circadian characteristics (n= 23 control, n= 17 antibiotic treatment): **A**, period; (z = -0.11, *p* = 0.912; Generalized Linear Mixed), values and error bars mean ± s.e.m., **B**, rhythm strength; (z = 0.243, *p* = 0.808, Generalized Linear Mixed Model), values and error bars mean ± s.e.m., **C**, Antibiotic treatment did not alter the mortality rate in honey bees. (χ2 = 0.08777, P = 0.7670, Log-Rank).

### Interaction with Nurse Bees

**Figure 2:**
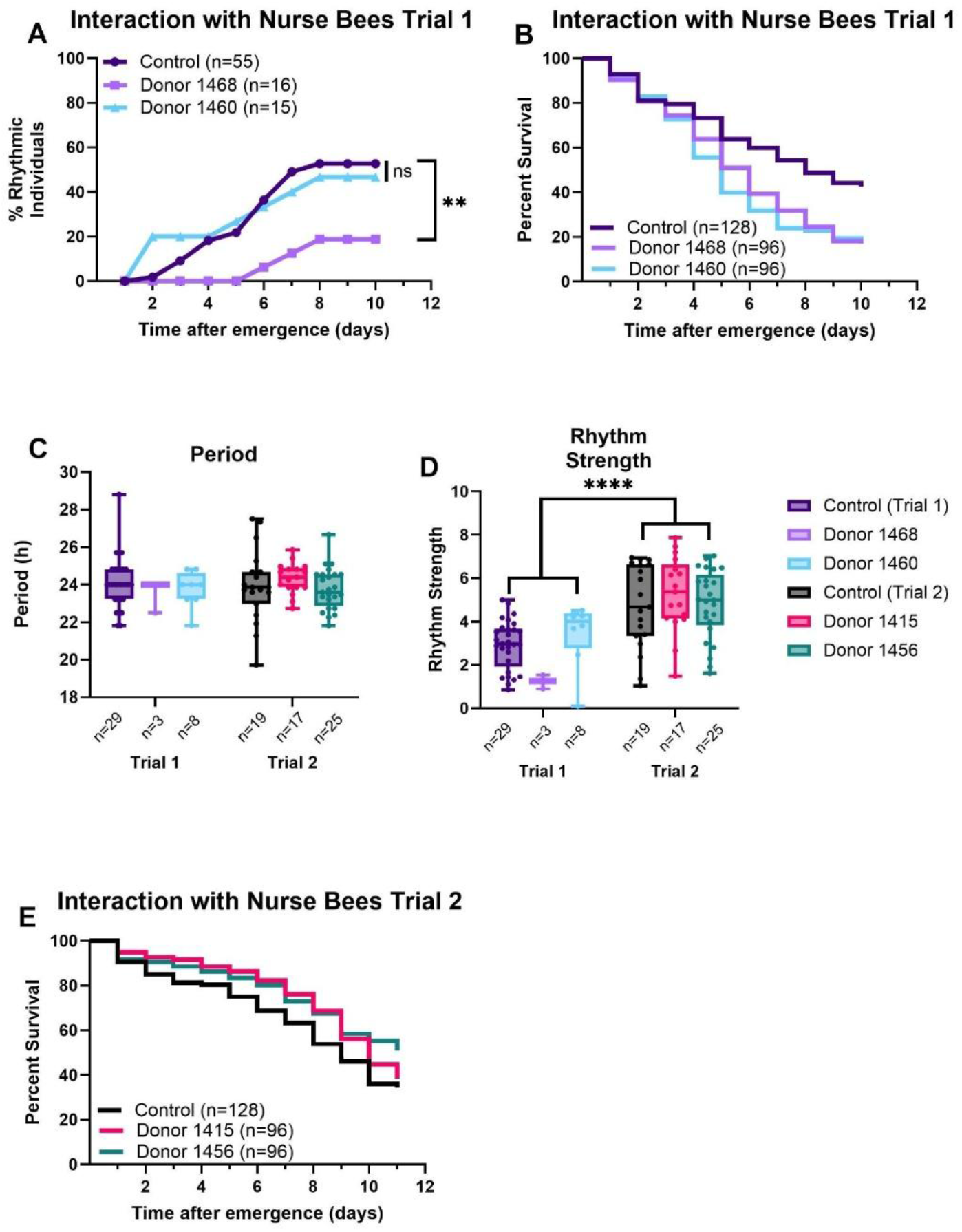
**A**, In the Trial 1, the donor 1468 group shows a lower percent of rhythmic individuals compared to other groups (n=55 control, n=16 donor 1468, n=15 donor 1460) Donor 1468: Wald = 6.65, p = 0.0099; Donor 1460: Wald = 0.17, p= 0.6870, Generalized Estimated Equations. **B**, the control group had a higher survival rate compared to the other two groups, which were newly-emerged bees that interacted with nurse bees (n=128 control, n=96 donor 1468, n=96 donor 1460) (χ^2^ = 22.42, P < 0.0001, Log-Rank). Two circadian parameters, periods and rhythm strength are examined using a two-way ANOVA (n=29 trial 1 control, n=3 donor 1468, n=8 donor 1460; n=19 control trial 2, n=17 donor 1415, n=25 donor 1456). **C**, shows period there were no significant differences (1460-Nurse Bees: z = 0.56, p = 0.58; 1468-Nurse Bees: z = −1.69, p = 0.09; Donor 1415: z = 0.67, p = 0.50; Donor 1456: z = −0.15, p = 0.88, Generalized Linear Mixed Model)), values and error bars mean ± s.e.m. **D**, exhibits rhythm strength; there are no significant differences (1460-Nurse Bees: z = 1.12, p = 0.26; 1468-Nurse Bees: z = −1.71, p = 0.087; Donor 1415: z = 1.29, p = 0.20; Donor 1456: z = 0.42, p = 0.67, Generalized Linear Mixed Model), values and error bars mean ± s.e.m. **E**, In the experiment, the control group had a lower survival rate than the groups from donor 1414 and donor 1456 (n=128 control, n=96 donor 1415, n=96 donor 1456) (χ2 = 6.772, P = 0.0339, Log-Rank).

### Brood Cap Experiments

**Figure 3:**
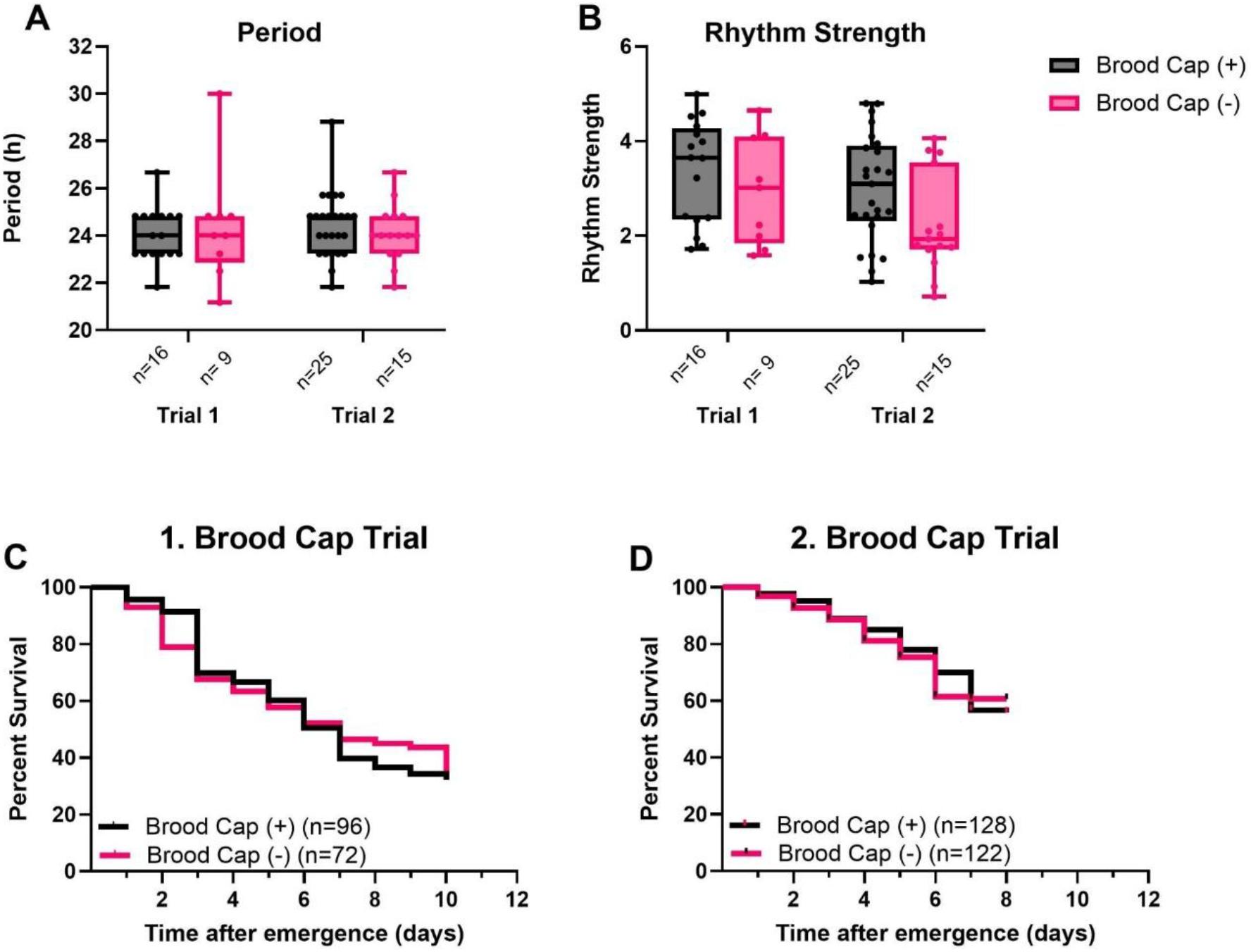
**A**, Period; no interaction between the trials and groups. No significant differences in the trials. Also, The treatment effect is not statistically significant (Brood Cap (+) vs Brood Cap (-); z = 0.95, p = 0.345, Generalized Linear Mixed Model), values and error bars mean ± s.e.m. **B**, Rhythm strength: no interaction was found between the trials and treatment. Additionally, there are no significant differences in both trials and treatment (Brood Cap (+) vs Brood Cap (-); z = 0.94, p = 0.346, Generalized Linear Mixed Model), values and error bars mean ± s.e.m. **C-D**) There were no differences in the mortality rate between the groups in both trials (χ2=0.005, P= 0.9436; χ2= 0.1432, P= 0.7051, Log-Rank).

**Figure 4:**
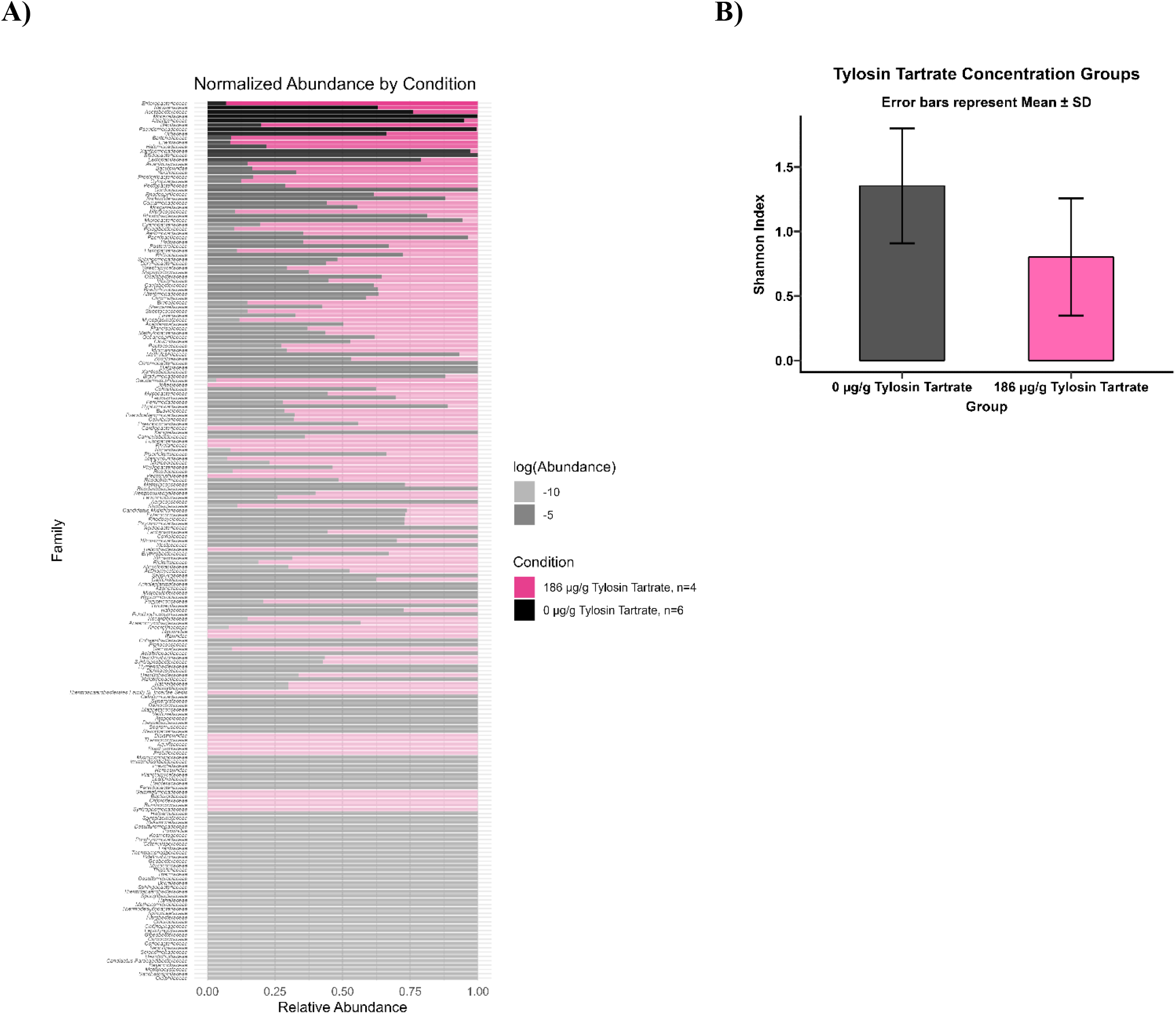
**A)** Antibiotic treatment alters gut microbial composition on genus level (F (1,9) = 1.8127, P = 0.0478, PERMANOVA). **B)** There were no statistical differences between control and treatment group (t = -1.904, P = 0.093, Generalized Linear Model, values and error bars mean ± s.d).

### Gut microbiota changed the brain transcriptomic profile in honey bees

**List 1.**
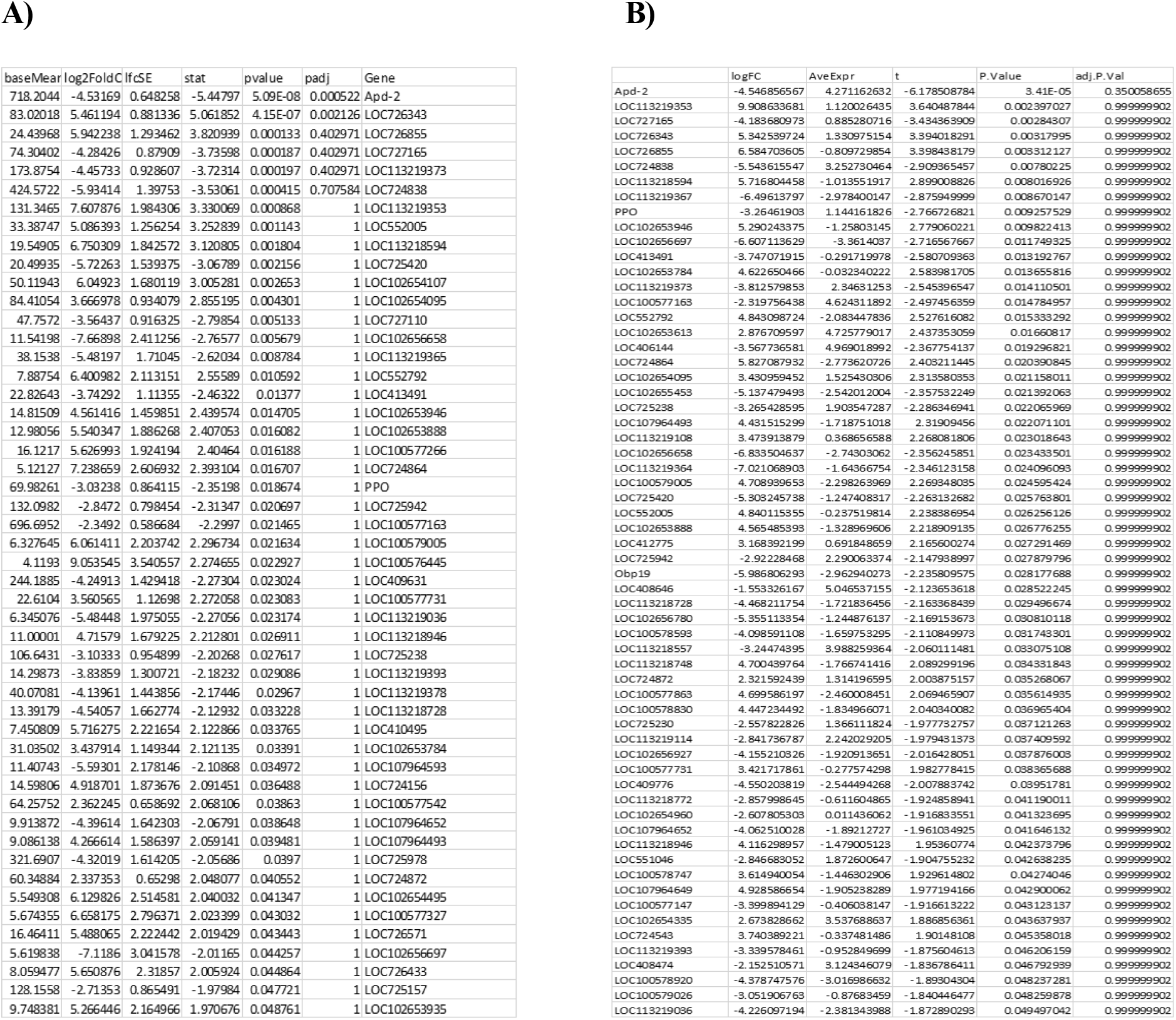
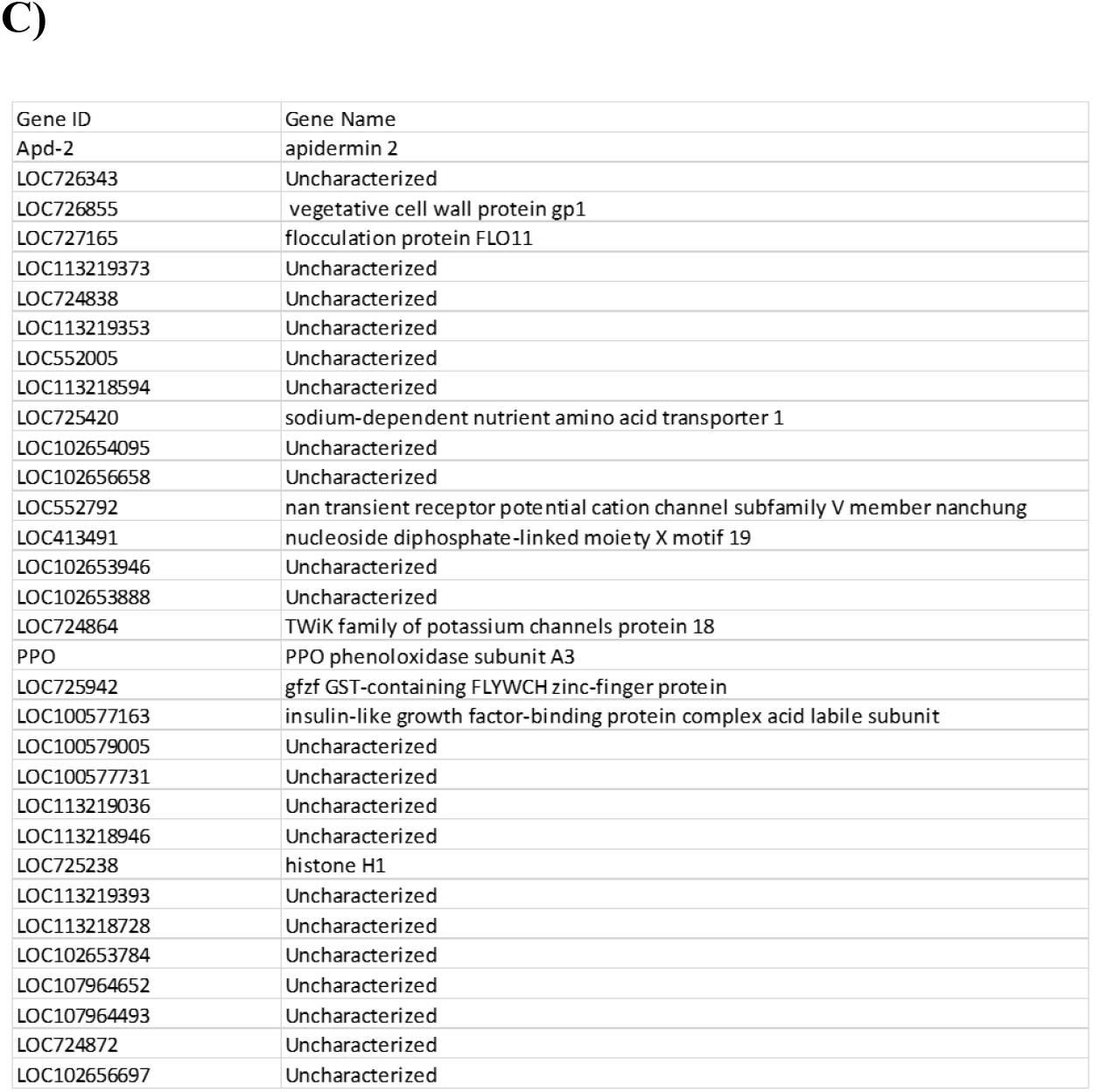
: Differentially expressed genes identified using two packages: **A)** DESeq2, **B)** limma, and **C)** genes common to both packages.

## Supplementary Discussion

Our results show that dysbiosis in the gut microbiota impairs the development of circadian rhythms and core clock mechanisms. We examined two key circadian parameters; period and rhythm strength in behaviorally rhythmic bees, alongside survival analyses from each behavioral experiment.

In the antibiotic treatment experiment, there were no significant differences between groups for either circadian parameter (Supplementary Fig. 1A–B). Moreover, survival was unaffected by antibiotic treatment (Supplementary Fig. 1C), suggesting that the treatment did not impair longevity under our experimental conditions.

For the second manipulation; nurse bee interaction, we conducted two separate trials. In Trial 1 (Supplementary Fig. 2A), differences emerged among donor groups: specifically, bees from Donor 1468 showed lower proportion of rhythmic individuals. Survival analysis revealed that the control group had a higher survival percentage than either donor group (Supplementary Fig. 2B). These results may reflect increased stress from aggressive interactions between nurse bees and newly emerged individuals. The limited sample size in donor groups may also have constrained our statistical power. Moreover, the stress experienced by newly emerged bees that interacted with donor groups was associated with reduced survival. However, the impact on circadian rhythm development may depend on the intensity of stress or colony-specific factors, as donor 1468 bees showed a markedly lower proportion of rhythmic individuals compared to donor 1460. In Trial 2, aggression was eliminated by using nurse bees from different colonies, resulting in similar proportions of rhythmic individuals across all groups and no significant differences (Fig. 1C). Interestingly, survival patterns in Trial 2 differed from those in Trial 1. Both donor groups showed significantly higher survival rates compared to the control group (Supplementary Fig. 2E). GLMM on circadian parameters revealed no significant effects on both period and rhythm strength.

As a third approach, we performed brood cap deprivation. We again used GLMM A to assess circadian parameters across treatment and experimental trial. No significant differences were observed for either period or rhythm strength (Supplementary Fig. 3A–B). Similarly, survival analysis from both trials showed no significant differences among groups (Supplementary Fig. 3C–D).

Taken together, these results indicate that gut microbiota does not significantly alter circadian parameters once rhythmicity is established. Locomotor rhythm characteristics remained consistent across groups, suggesting that period and rhythm strength are regulated independently of the gut microbiota. This aligns with findings in *Drosophila*, where Silva et al. (2021) showed that microbiota do not affect periodicity. Additionally, we found that neither gut microbiota depletion by antibiotics nor brood cap depletion impaired honey bee survival. This contrasts with Raymann et al. (2017), who observed reduced lifespan in bees treated with higher antibiotic dosages. Our study used 186 µg/g, whereas Raymann et al. applied 450 µg/mL which is approximately 2.3 times more indicating a potential dosage-dependent effect on mortality. Also, Liberti et al 2022 show no difference in survival between microbiota-deprived and conventional bees.

To assess microbial composition, we performed rRNA-depleted RNA sequencing on gut tissue and profiled bacterial taxa using Kraken2. Significant shifts at the family level were observed following antibiotic treatment. Core bacterial families decreased in response to antibiotic treatment, including Neisseriaceae, Bifidobacteriaceae, Lactobacillaceae, and Orbaceae. These families include key members of the honey bee core microbiota, such as *Snodgrassella, Bifidobacterium, Lactobacillus*, and *Gilliamella*, respectively (Motta & Moran, 2024).

Previous work supports the role of gut microbiota in shaping host behavior and gene expression in the brain. For instance, Zheng et al. (2017) reported that gut bacteria regulate *insulin-like peptide 1/2* expression in the honey bee head. Liberti et al. (2022) found that microbe-deprived bees engage in fewer head-to-head interactions, and that *Bifidobacterium asteroides* affect many brain gene expression changes observed in bees colonized with a full microbiota. Similarly, Zhang et al. (2022) demonstrated that *Lactobacillus Firm-5* enhances learning and memory, and that different bacterial taxa such as *Gilliamella* and *Lactobacillus Firm4/Firm5* exert distinct effects on brain gene expression, metabolite profiles, and neurotransmitter levels. Notably, mammals share core bacterial genera with honey bees including *Lactobacillus* and *Bifidobacterium*. Exposure to Lactobacillus and *Bifidobacterium* in the fetal and early postnatal ages promotes neural development and behavior including learning, social interaction, anxiety-like behavior^17–20^. Also our data suggest that the microbes from the brood cap, to which honey bees are potentially first introduced, may affect the development of circadian rhythm. Our study showed for the first time that disruption of gut microbiota during early postnatal ages impaired the development of circadian rhythm and clock system in honey bees.

## Notes

### Competing Interest Statement

The authors have declared no competing interest.

### Summary of Updates

The analysis found in the Figure 2 and Supplementary Figure 4 A and B updated. Also, graphs are fixed

### Funding

- NIH: 5P20GM103642, 2R25NS080687, P20GM103475
- NSF: 1736026, 1633184, 2131647, Arecibo Ac3
- USDA: 2021-67014-34999
- Volkswagen Foundation
- NLM: 2T15LM007059-36
- Deutsche Forschungsgemeinschaft (DFG, German Research Foundation): (BE8339/1-1)

## REFERENCES

1. Jabbari Shiadeh, S.M., Chan, W.K., Rasmusson, S. et al. Bidirectional crosstalk between the gut microbiota and cellular compartments of brain: Implications for neurodevelopmental and neuropsychiatric disorders. Transl Psychiatry 15, 278 (2025).

2. Comas, M., De Pietri Tonelli, D., Berdondini, L. & Astiz, M. Ontogeny of the circadian system: a multiscale process throughout development. Trends Neurosci. 47, 36–46 (2024).

3. Moore, D., Angel, J. E., Cheeseman, I. M., Fahrbach, S. E. & Robinson, G. E. Timekeeping in the honey bee colony: Integration of circadian rhythms and division of labor. Behav. Ecol. Sociobiol. 43, 147–160 (1998).

4. Eban-Rothschild, A., Shemesh, Y. & Bloch, G. The colony environment, but not direct contact with conspecifics, influences the development of circadian rhythms in honey bees. J. Biol. Rhythms 27, 217–225 (2012).

5. Beer, K. & Helfrich-Förster, C. Post-embryonic Development of the Circadian Clock Seems to Correlate With Social Life Style in Bees. Front. Cell Dev. Biol. 8, 1–9 (2020).

6. Rivkees, S. A. Developing circadian rhythmicity in infants. Pediatr. Endocrinol. Rev. 1, 38–45 (2003).

7. Baxter, R. C. Endocrine and cellular physiology and pathology of the insulin-like growth factor acid-labile subunit. Nat. Rev. Endocrinol. 20, 414–425 (2024).

8. Robinson, H., Dave, N., Barzilay, R. et al. The effect of the “exposome” on developmental brain health and cognitive outcomes. Neuropsychopharmacol. (2025).

9. Landgral, D., Koch, C. E. & Oster, H. Embryonic development of circadian clocks in the mammalian suprachiasmatic nuclei. Front. Neuroanat. 8, 1–7 (2014).

10. Beer, K. et al. Pigment-Dispersing Factor-expressing neurons convey circadian information in the honey bee brain. Open Biol. 8, (2018).

11. Fagiani, F. et al. Molecular regulations of circadian rhythm and implications for physiology and diseases. Signal Transduct. Target. Ther. 7, 1–20 (2022).

12. Zheng, X., Zhang, K., Zhao, Y. & Fent, K. Environmental chemicals affect circadian rhythms: An underexplored effect influencing health and fitness in animals and humans. Environ. Int. 149, 106159 (2021).

13. Fishbein, A. B., Knutson, K. L. & Zee, P. C. Circadian disruption and human health. J. Clin. Invest. 131, (2021).

14. Walker, W. H., Walton, J. C., DeVries, A. C. & Nelson, R. J. Circadian rhythm disruption and mental health. Transl. Psychiatry 10, (2020).

15. Brown, M. R. et al. Time-restricted feeding prevents deleterious metabolic effects of circadian disruption through epigenetic control of β cell function. Sci. Adv. 7, 1–19 (2021).

16. Leone, V. et al. Effects of diurnal variation of gut microbes and high-fat feeding on host circadian clock function and metabolism. Cell Host Microbe 17, 681–689 (2015).

17. Thaiss, C. A. et al. Microbiota Diurnal Rhythmicity Programs Host Transcriptome Oscillations. Cell 167, 1495–1510.e12 (2016).

18. Tofani, G. S. S. et al. Gut microbiota regulates stress responsivity via the circadian system. Cell Metab. 37, 138–153.e5 (2024).

19. Jin, Y., Wu, S., Zeng, Z. & Fu, Z. Effects of environmental pollutants on gut microbiota. Environ. Pollut. 222, 1–9 (2017).

20. Lynch, C. M. K. et al. Critical windows of early-life microbiota disruption on behaviour, neuroimmune function, and neurodevelopment. Brain. Behav. Immun. 108, 309–327 (2023).

21. Luck, B. et al. Bifidobacteria shape host neural circuits during postnatal development by promoting synapse formation and microglial function. Sci. Rep. 10, 1–18 (2020).

22. Volkova, A. et al. Effects of early-life penicillin exposure on the gut microbiome and frontal cortex and amygdala gene expression. iScience 24, 102797 (2021).

23. Li, J. et al. Antibiotic cocktail-induced gut microbiota depletion in different stages could cause host cognitive impairment and emotional disorders in adulthood in different manners. Neurobiol. Dis. 170, 105757 (2022).

24. O’Connor, R. et al. Maternal antibiotic administration during a critical developmental window has enduring neurobehavioural effects in offspring mice. Behav. Brain Res. 404, (2021).

25. Heijtz, R. D. et al. Normal gut microbiota modulates brain development and behavior. Proc. Natl. Acad. Sci. U. S. A. 108, 3047–3052 (2011).

26. Erny, D. et al. Host microbiota constantly control maturation and function of microglia in the CNS. Nat. Neurosci. 18, 965–977 (2015).

27. Lu, J. et al. Effects of Intestinal Microbiota on Brain Development in Humanized Gnotobiotic Mice. Sci. Rep. 8, 1–16 (2018).

28. Zheng, H., Steele, M. I., Leonard, S. P., Motta, E. V. S. & Moran, N. A. Honey bees as models for gut microbiota research. Lab Anim. (NY). 47, 317–325 (2018).

29. Bloch, G., Toma, D. P. & Robinson, G. E. Behavioral rhythmicity, age, division of labor and period expression in the honey bee brain. J. Biol. Rhythms 16, 444–456 (2001).

30. Beer, K., Steffan-Dewenter, I., Härtel, S. & Helfrich-Förster, C. A new device for monitoring individual activity rhythms of honey bees reveals critical effects of the social environment on behavior. J. Comp. Physiol. A Neuroethol. Sensory, Neural, Behav. Physiol. 202, 555–565 (2016).

31. Giannoni-Guzmán, M. A. et al. The role of temperature on the development of circadian rhythms in honey bee workers. PeerJ 12, 1–15 (2024).

32. Fahrbach, S. E. & Robinson, G. E. Behavioral development in the honey bee: toward the study of learning under natural conditions. Learn. Mem. 2, 199–224 (1995).

33. Fahrbach, S.E. & Robinson, G.E. Juvenile Hormone, Behavioral Maturation, and Brain Structure in the Honey Bee. Dev Neurosci. 18, 102–114 (1996).

34. Seeley, T. D. Adaptive significance of the age polyethism schedule in honeybee colonies. Behav. Ecol. Sociobiol. 11, 287–293 (1982).

35. Ortiz-Alvarado, Y. et al. Antibiotics in hives and their effects on honey bee physiology and behavioral development. Biol. Open 9, (2020).

36. Bloch, G., Sullivan, J. P. & Robinson, G. E. Juvenile hormone and circadian locomotor activity in the honey bee Apis mellifera. J. Insect Physiol. 48, 1123–1131 (2002).

37. Champagne-Jorgensen, K., Kunze, W. A., Forsythe, P., Bienenstock, J. & McVey Neufeld, K. A. Antibiotics and the nervous system: More than just the microbes? Brain. Behav. Immun. 77, 7–15 (2019).

38. Martinson, V. G., Moy, J. & Moran, N. A. Establishment of characteristic gut bacteria during development of the honeybee worker. Appl. Environ. Microbiol. 78, 2830–2840 (2012).

39. Powell, J. E., Martinson, V. G., Urban-Mead, K. & Moran, N. A. Routes of acquisition of the gut microbiota of the honey bee Apis mellifera. Appl. Environ. Microbiol. 80, 7378–7387 (2014).

40. Motta, E. V. S. & Moran, N. A. The honeybee microbiota and its impact on health and disease. Nature Reviews Microbiology vol. 22 (2024).

41. Fuchikawa, T. et al. Neuronal circadian clock protein oscillations are similar in behaviourally rhythmic forager honeybees and in arrhythmic nurses. Open Biol. 7, (2017).

42. Fernandez, A. M. & Torres-Alemán, I. The many faces of insulin-like peptide signalling in the brain. Nat. Rev. Neurosci. 13, 225–239 (2012).

43. Giannoni-Guzmán, M. A. et al. Measuring individual locomotor rhythms in honey bees, paper wasps and other similar-sized insects. J. Exp. Biol. 217, 1307–1315 (2014).

## References

1. Giannoni-Guzmán, M. A. et al. Measuring individual locomotor rhythms in honey bees, paper wasps and other similar-sized insects. J. Exp. Biol. 217, 1307–1315 (2014).

2. Giannoni-Guzmán, M. A. et al. The Role of Temperature on the Development of Circadian Rhythms in Honey Bee Workers. BioRxiv (2020) doi:10.1101/2020.08.17.254557.

3. Powell, J. E., Martinson, V. G., Urban-Mead, K. & Moran, N. A. Routes of acquisition of the gut microbiota of the honey bee Apis mellifera. Appl. Environ. Microbiol. 80, 7378–7387 (2014).

4. Zhang, Z. et al. Honeybee gut Lactobacillus modulates host learning and memory behaviors via regulating tryptophan. Nat. Commun. 13, 1–13 (2022).

5. Martinson, V. G., Moy, J. & Moran, N. A. Establishment of characteristic gut bacteria during development of the honeybee worker. Appl. Environ. Microbiol. 78, 2830–2840 (2012).

6. Beer, K. & Helfrich-Förster, C. Post-embryonic Development of the Circadian Clock Seems to Correlate With Social Life Style in Bees. Front. Cell Dev. Biol. 8, 1–9 (2020).

7. Pende, M. et al. A versatile depigmentation, clearing, and labeling method for exploring nervous system diversity. Sci. Adv. 6, (2020).

8. Fuchikawa, T. et al. Neuronal circadian clock protein oscillations are similar in behaviourally rhythmic forager honeybees and in arrhythmic nurses. Open Biol. 7, (2017).

9. Beer, K. et al. Pigment-Dispersing Factor-expressing neurons convey circadian information in the honey bee brain. Open Biol. 8, (2018).

10. Bolger, A. M., Lohse, M. & Usadel, B. Trimmomatic: A flexible trimmer for Illumina sequence data. Bioinformatics 30, 2114–2120 (2014).

11. Dobin, A. et al. STAR: Ultrafast universal RNA-seq aligner. Bioinformatics 29, 15–21 (2013).

12. Giannoni-Guzmán, M. A. et al. The role of temperature on the development of circadian rhythms in honey bee workers. PeerJ 12, 1–15 (2024).

13. Silva, V. et al. The impact of the gut microbiome on memory and sleep in Drosophila. J. Exp. Biol. 224, (2021).

14. Raymann, K., Shaffer, Z. & Moran, N. A. Antibiotic exposure perturbs the gut microbiota and elevates mortality in honeybees. PLoS Biol. 15, 1–22 (2017).

15. Liberti, J. et al. The gut microbiota affects the social network of honeybees. Nat. Ecol. Evol. 6, 1471–1479 (2022).

16. Zheng, H., Powell, J. E., Steele, M. I., Dietrich, C. & Moran, N. A. Honeybee gut microbiota promotes host weight gain via bacterial metabolism and hormonal signaling. 114, 4775–4780 (2017).

17. Luck, B. et al. Bifidobacteria shape host neural circuits during postnatal development by promoting synapse formation and microglial function. Sci. Rep. 10, 1–18 (2020).

18. Luk, B. et al. Postnatal colonization with human ‘infant-type’ Bifidobacterium species alters behavior of adult gnotobiotic mice. PLoS One 13, 1–25 (2018).

19. Lopez-Tello, J. et al. Maternal gut Bifidobacterium breve modifies fetal brain metabolism in germ-free mice. Mol. Metab. 88, 1–12 (2024).

20. Buffington, S. A. et al. Microbial Reconstitution Reverses Maternal Diet-Induced Social and Synaptic Deficits in Offspring. Cell 165, 1762–1775 (2016).

